# Major Contribution of Myeloid Cells In TB specific Host Gene Signature: Revelations from Re-Analysis of Publicly Available Datasets

**DOI:** 10.1101/2022.09.27.508979

**Authors:** Anuradha Gautam, Saroj Kant Mohapatra, Bhaswati Pandit

## Abstract

**Introduction:** Interaction of human host and its pathogen *M.tuberculosis* drives tuberculosis disease, resulting in dysregulation of host gene expression. We re-analyzed host gene expression datasets of TB to identify and validate a cellular circuit by interlinking the DEGs with their target miRNAs, GWAS hits associated with immunological phenotypes mapping to the DEGs and associated cellular subtypes through bioinformatic and experimental approaches.

**Methodology:** DEGs were identified systematically through re-analysis of whole blood host transcriptomic datasets of treatment-naive TB patients, obtained from public repositories having at least 1.2 fold change of expression and FDR corrected p-value <0.05. Using well characterized *M.tb* antigens: Ag85 complex, LAM, CFP10 and ESAT6, we evaluated their effect on the expression of a subset of the top DEGs with at least two fold change of expression in a monocytic cell line THP1 with or without differentiation into a macrophage-like state with PMA and a T cell line Jurkat E6-1.

**Results:** We discovered 305 DEGs (236 up and 69 down-regulated genes) out of which 23 (21 up and 2 down-regulated genes) were top DEGs. Overall, innate immune and myeloid cell associated pathways were enriched for up-regulated genes while T cell associated pathways were enriched for down-regulated genes. Among top DEGs, *EPSTI1* was predominantly up-regulated in macrophages while *SERPING1* was universally up-regulated in the monocyte model by all antigens. The down-regulation of gene expression was replicated by ESAT6 in T cell line by significantly down-regulating a top down-regulated gene *LRRN3*.

## Introduction

Host gene expression in tuberculosis has emerged as a tool to identify immune signatures associated with the disease. These signatures find application in disease diagnosis, exploration of disease progression and response to therapy. Dissection of host whole blood transcriptome using microarray and high-throughput sequencing platforms have helped identify gene signatures to diagnose TB with [1–5]□. An early whole blood transcriptome study showed IFNγ and IFNα/β signalling circuits to be driven by neutrophils in tuberculosis [1]□. Interferon-inducible signatures were observed to be similar in granulomatous diseases like pulmonary TB and sarcoidosis. However, the abundance and intensity of expression of these transcripts was greater in TB compared to sarcoidosis [3]□. Further, the study described a 144 transcript signature which provided specificity of TB detection from other lung diseases [3]□. Transcriptomic signatures also vary between pulmonary and latent tuberculosis. A 380 metagene signature was observed to be specific towards the detection of pulmonary TB while showing heterogeneity of expression in LTBI [4] □. The expression of *DOCK9*, *EPHA4* and *NPC2* transcripts were observed to be up-regulated in TB patients from Brazil compared to exposed controls [5]□. As a guanine exchange factor (GEF), DOCK9 activated CDC42. EPHA4 is a protein tyrosine kinase and is associated with GPCR pathway and ERK signalling pathways. NPC2 is a cholesterol transport regulator for the late endosomal/lysosomal system. These findings were later validated by RT-PCR and re-analysis of gene expression data from different areas of diverse genetic background: South Africa, Germany and France.

A diagnostic transcriptomic signature is important in paediatric tuberculosis where detection of TB is difficult. A 51-gene signature was discovered in cohorts of Malawian and South African children and validated in Kenyan children distinguishing TB from other diseases, irrespective of HIV co-infection [6]□. This host-derived gene signature was comparatively more sensitive than commercially available Xpert MTB/RIF assay in diagnosing culture negative TB cases among Kenyan children [6]□. In another analysis involving geographically distinct cohorts from Uganda and India, a two-genes *RAB20* and *INSL3* could successfully diagnose TB in PLWH [7]□.

In addition to demonstrating specificity of TB detection, host gene expression profiles also capture and characterize the clinical stages of tuberculosis and response to therapy. A 15-transcript signature was observed distinguishing active TB cases from latent TB cases (TST+) and unexposed controls in the absense of HIV co-infection [8]□. Advances in single cell sequencing have helped decode contribution of different cell types in the process of infection. The composition of human PBMCs has been shown to change in active and latent tuberculosis. This change was significant in the subsets of NK cells which varied in prevalence in active TB, LTBI and healthy controls [9]□. Although sputum conversion is a surrogate for effective TB treatment, it has been observed that a whole blood IFN-dominanted transcriptome signature could also be used to monitor therapy using an anti-tuberculosis therapy (ATT) specific 320-transcript signature [10]□.

In our study, we have re-analyzed publicly available datasets of host whole blood transcriptome in treatment-naive TB cases for the identification of DEGs and their functional validation in a cellular model. Whole blood transcriptomic studies are important as they retain gene expression signals from granulocytes too which would have otherwise been lost from PBMC datasets. This re-analysis revealed an enrichment of inflammatory pathways, innate immune pathways and myeloid leukocyte activation pathways among the up-regulated genes. Adaptive immunity and T cell associated pathways were enriched within the down-regulated genes. Genes with two fold change in expression were identified as top DEGs which showed activatory marks in CD14^+^ monocytes and neutrophils. Notably, many of the top up-regulated genes were a part of the “response to interferon” pathway of GO, biological processes. Indicating an overlap of host genomic factors onto the dynamic changes in the host transcriptome during TB disease, GWAS significant SNPs associated with phenotypes like CRP level, IFNγ and MCSF levels mapped to top DEGs *AIM2, DHRS9* and *C1QB.*

A subset of the top DEGs were selected for functional validation in a model of THP1 monocytes/macrophages and T cell line Jurkat E6-1 treated with four TB antigens; Ag85 complex, LAM, CFP10 and ESAT6. In THP1 monocytes, LAM showed an elevated expression of genes *EPSTI*1 and *AIM*2 compared to the Ag85 complex and CFP10 treatment. ESAT6 treatment induced the greatest fold up-regulation of *SERPING*1 and *GBP5* expression in monocytes. In macrophages, ESAT6 treatment showed most up-regulation for the expression of *GBP*5 and *EPSTI*1. *EPSTI*1 was up-regulated by the Ag85 complex, LAM and ESAT6. Overall, *EPSTI1* overexpression was inducible by the Ag85 complex, LAM and ESAT6 in PMA derived macrophages while *SERPING*1 was significantly overexpressed by all four antigens in the monocytes. In the T cell line model, ESAT6 was observed to down-regulate the expression of *LRRN3* which was identified as a top down-regulated gene. These findings depicts the roles of different mycobacterial antigens and their contribution towards activating/inactivating differentially expressed genes in tuberculosis. Overview of the study design has been provided in Figure 1.

**Figure 1:**
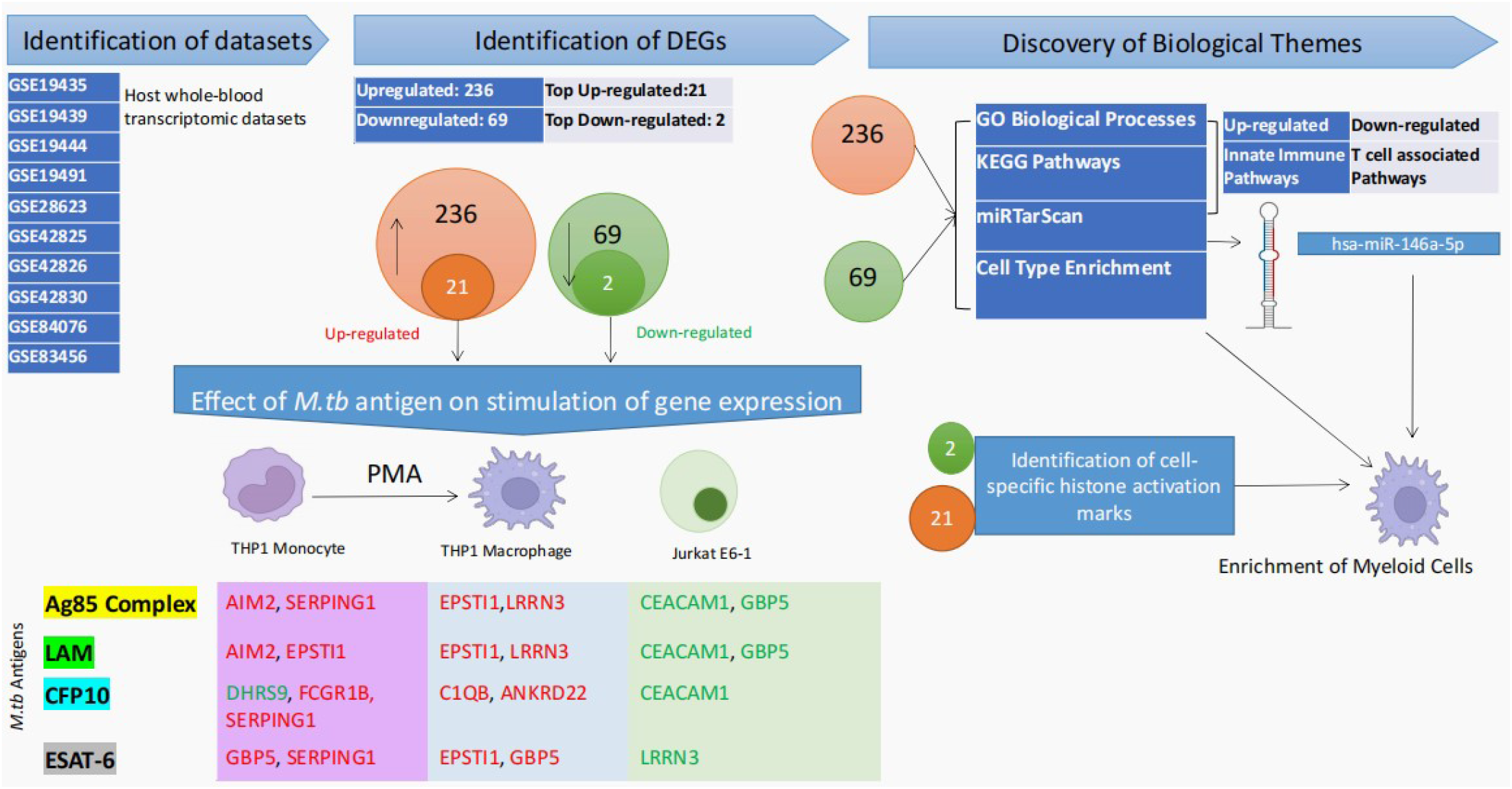
Overview of Study: Whole blood host transcriptomic datasets were queried from public repositories. Differentially expressed genes (DEGs) were identified and pathway analysis and other enrichment analyses were carried out to derive biologically relevant themes from the data. A subset of the top DEGs were used to study the effect of *M.tb* antigens on host gene expression in a monocyte-macrophage model and T cell model.

## Materials and Methods

### 1. Identification of Host transcriptomic datasets

Host whole blood TB transcriptomic datasets were retrieved from three databases: PubMed, Gene Expression Omnibus (GEO) and ArrayExpress using the search strings (tuberculosis OR “Mycobacterium tuberculosis” OR “pulmonary tuberculosis”) AND (“gene expression” OR “Host gene expression” OR “host transcriptome” OR ‘transcriptome”) on 27th November 2018. This search included datasets published from January 2001 (01-01-2001) to August 2018 (31-08-2018) across the three databases.

203 records were retrieved from GEO (143) and ArrayExpress (60). 1562 records were obtained from Pubmed. All records were manually sorted, cross-referenced and merged to recover 142 datasets. The curation of records and processes used to filter through these records have been described in Fig 2. Following filtering, 10 whole blood host transcriptomics studies comparing treatment-naive, HIV-negative pulmonary TB cases and healthy controls were selected for further analysis. The details of the studies retained and sample size after quality control have been mentioned in Table 1.

**Figure 2:**
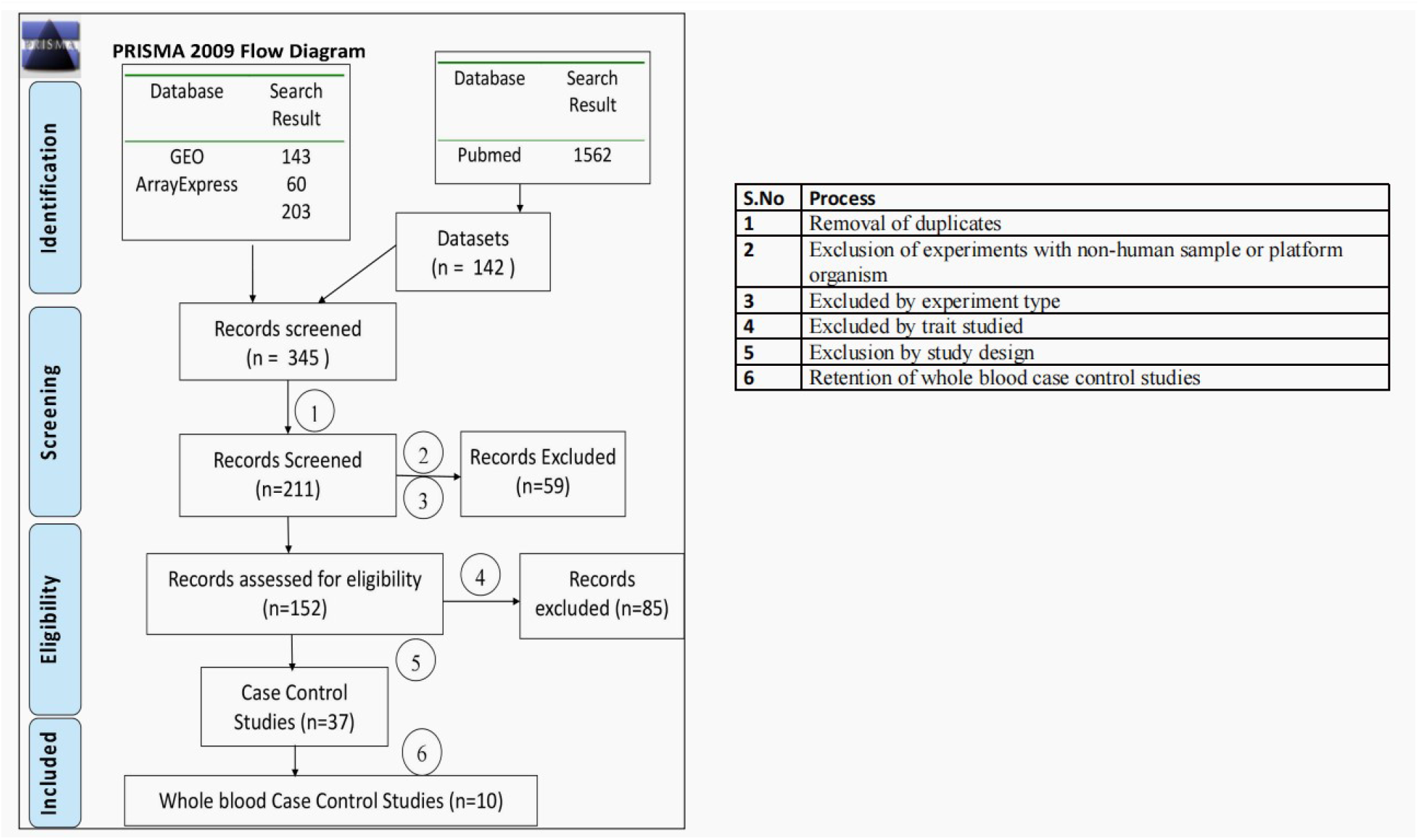
Identification of Datasets for Re-analysis: PRISMA Flow chart showing the identification and inclusion of records for the meta analysis. The numbers refer to the processes performed to include or exclude a study.

**Table 1:** Overview of Studies Included for re-analysis

| S. N | GEO Accession ID | Author, Year | Sample Size | Platform Technology |
| --- | --- | --- | --- | --- |
| 1 | GSE19435 | MP Berry, 2010 | 25 | Array |
| 2 | GSE19439 | MP Berry, 2010 | 33 | Array |
| 3 | GSE19444 | MP Berry, 2010 | 32 | Array |
| 4 | GSE19491 | MP Berry, 2010 | 155 | Array |
| 5 | GSE28623 | J Maertzdorf, 2011 | 79 | Array |
| 6 | GSE42825 | CI Bloom, 2013 | 62 | Array |
| 7 | GSE42826 | CI Bloom, 2013 | 31 | Array |
| 8 | GSE42830 | CI Bloom, 2013 | 54 | Array |
| 9 | GSE84076 | LS deAraujo, 2016 | 18 | HTS |
| 10 | GSE83456 | S Blankley, 2016 | 105 | Array |

### 2. Data download and processing

The series matrix files corresponding to each dataset were downloaded from GEO. For microarray studies, the probes without gene annotation were removed. Duplicated probes i.e., multiple probes corresponding to the same gene were filtered using the function nsFilter() of the R package genefilter to retain the probes with highest variance. The probe IDs were replaced with Entrez Gene names using org.Hs.eg.db [11]□. Quality control of samples was performed using arrayQuality metrics. Samples failing quality control were excluded from further analysis. Following quality control, a total of 594 samples were retained for further analyses. All datasets were harmonized by considering only the common genes present on all platforms.

### 3. Identification of Differentially Expressed Genes (DEGs)

Two-tailed, two-sample ‘t’ test was performed to identify the DEGs. Up-regulated and down-regulated genes were selected on the bases of fold change (at least 20% or 1.2 fold change in cases compared to the controls) and significance (FDR corrected p-value <0.05). The top DEGs were identified with a two fold change of expression.

### 4. Pathway Analysis

To explore the putative biological functions and pathways enriched for the DEGs, over-representation analysis (ORA) was performed. Gene Ontology (GO) and KEGG pathway assignment was carried out using clusterProfiler package [12]□. To aid interpretation of the enriched terms we used tree plots and enrichment maps to simplify and cluster the enriched terms. To explore their regulation, enrichment of miRNA targets for the DEGs was carried out using Enrichr https://maayanlab.cloud/Enrichr/ [13–15]□ and the significant DEG-miRNA interactions were visualized with miRNet2.0 [16]□.

### 5. Identification of active cellular subsets

The HuBMAP and Human Gene Atlas gene set libraries were used in Enrichr to identify cell types enriched for the up and down-regulated genes. HuBMAP library consists of cell type biomarkers curated by the HuBMAP consortium [17]□ while the Human Gene Atlas uses the BioGPS dataset [18]□ for enrichment of tissue specific genes. Further, presence of histone activatory marks around the promoters of the top DEGs were visualized from ENCODE [19]□ ChIP-seq tracks in UCSC genome browser (hg19). Promoter coordinates for the top DEGs were found from EPD [20]□ and GeneHancer [21]□ tracks in UCSC genome browser. The presence or absence of histone activatory marks around the promoter was visualized in Microsoft Excel for each cell type.

### 6. Assessing the effect of *M.tb* antigens on gene expression in monocytic, macrophage like cells and a T cell line

Four *M.tb* antigens: Ag85 complex, LAM, CFP10, and ESAT6 were selected to test for the induction of gene expression in a model of THP1 monocytes. The antigens were sourced from BEI Resources. To insinuate a monocytic model, THP1 cells were directly stimulated with 5μg/ml of CFP10, LAM, Ag85 or ESAT6 for 24 hours. Similarly, T cell line Jurkat E6-1 was also directly treated with 5ug/ml of the above mentioned anitgens for a period of 24 hours. Monocytic THP1 were treated with 100nM PMA for 24h to undergo differentiation and the resultant macrophage like cells were treated similarly with the *M.tb* antigens.

RNA was isolated using TRIzol reagent and treated with DNase I. cDNA synthesis was carried with RNA input of 500-1000 μg using Verso cDNA synthesis kit and RT-PCR was performed. The Ct values for each condition was normalized against B-actin expression for the monocytes/marophage like cells and against GAPDH for the T cell line. The fold change was calculated for monocytic and T cell model against untreated THP1 and Jurkat E6-1 respectively. In the macrophages, the fold change was normalized against PMA treated THP1 for the macrophage model. Each antigenic stimulation was performed in duplicates and RT-PCR readings were taken in duplicates. A Ct value cutoff was selected at 32, any sample or gene with a Ct value greater than 32 was considered to be undetectable expression. List of primers for RT-PCR is provided in supplementary Table 4

## Results

### 1. Identification of DEGs

DEGs with 20% or 1.2 fold change in expression and FDR corrected p < 0.05 were identified by performing two sample two tailed t test using the function *rowttests* in R. These DEGs include 236 up-regulated genes and 69 down-regulated genes. Top DEGs were identified with having a 2 fold change of expression and FDR corrected p value <0.05. A subset of top DEGs were selected for studying the effect of *M.tb* antigens on their expression in a monocyte/macrophage and T cell model by RT-PCR. GO, KEGG and miRTarScan over representation analysis was carried out on these DEGs (236 up-regulated and 69 down-regulated genes) to identify enriched biological pathways and miRNA targets within the DEGs.

### 2. Pathway analysis of Up-regulated genes

Enrichment analysis of GO Biological Processes and KEGG pathways was carried out on the 236 up-regulated genes. Top enriched GO Biological Processes include *IFN-γ response pathways* (“response to interferon gamma” and “cellular response to interferon-gamma”) and *innate immune pathways* (“regulation of innate response”, “pattern recognition receptor”, “myeloid cell activation”, “toll like receptor signalling”, and “regulation of inflammatory response”) (Fig 3A). Clustering all enriched terms revealed five clusters; *I-kappa inflammatory kinase/ NF-kappa B interferon gamma, pattern recognition response regulating receptor, positive innate biotic external, defense to virus symbiont and regulation of interleukin-1 beta formation* (Fig 3C). The relevant TB associated pathways which include the IFN-γ response pathways, and myeloid leukocyte associated pathways are a part of the I-kappa inflammatory kinase/ NF-kappa B interferon gamma cluster. Enrichment maps help visualize interactions between mutually overlapping gene sets or pathways which cluster together. Enriched GO Biological Processes showed distinct clusters associated with interferon γ response, virus response, response to biotic stimulus and pattern recognition receptor signaling (Fig 3E).

**Figure 3:**
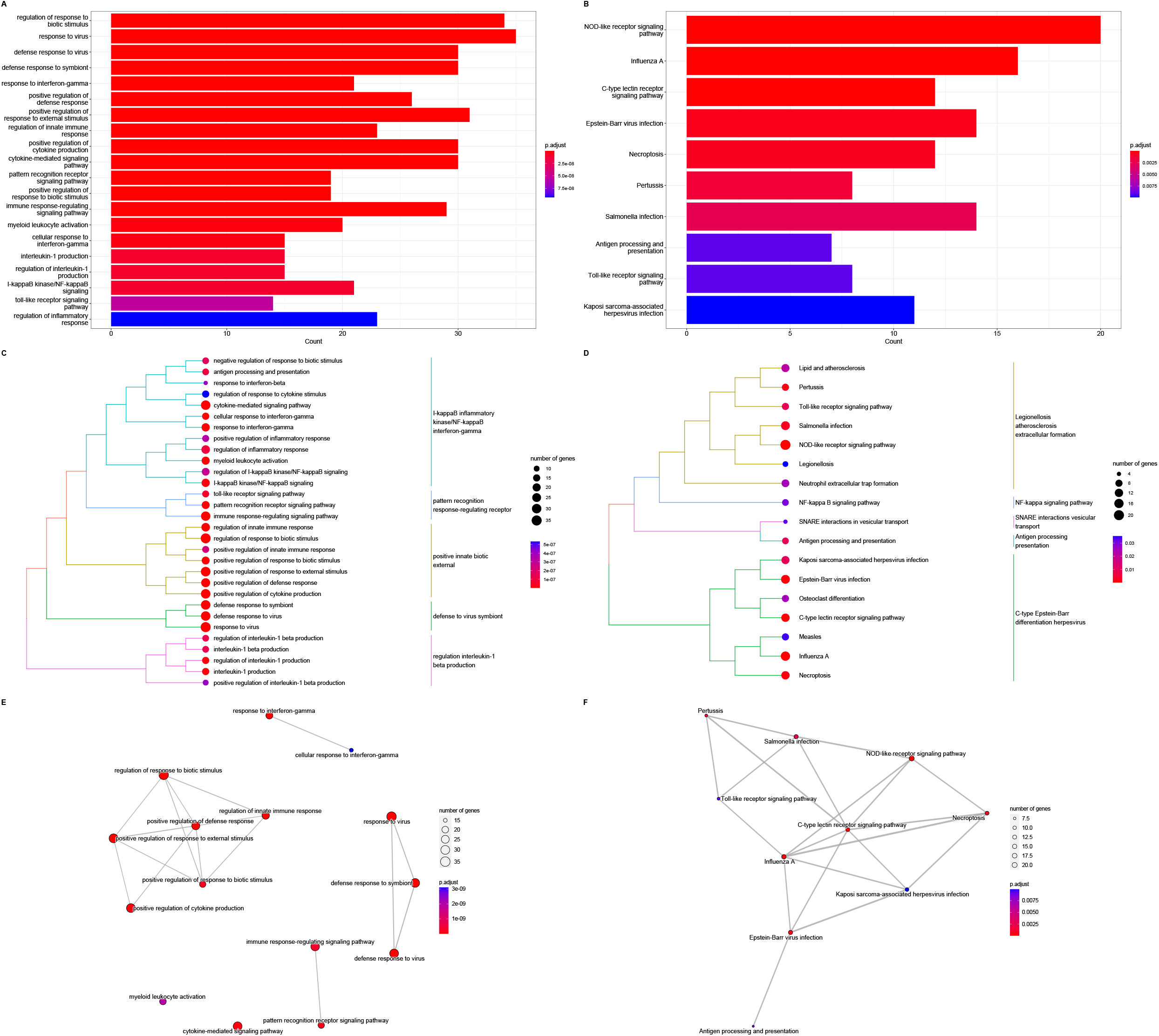
Pathway Analysis for Up-regulated Genes using GO and KEGG pathways: Bar plot shows enriched biological pathways for the up-regulated genes GO Biological Processes (3A) and KEGG pathways (3B). To simplify the enriched terms, tree plot was made to identify types of pathways enriched within the up-regulated genes for GO Biological Processes (3C) and KEGG pathways (3D). Enrichment map of GO Biological Processes shows four distinct clusters (3E). while KEGG pathways show interconnected pathways (3F).

Top enriched KEGG pathways also featured innate immune processes including recognition through PRR like “nod-like receptor signalling”, “c-type lectin receptor signalling pathway” and “toll-like receptor signalling pathway” (Fig 3B). Other innate immune pathways include “necroptosis” and “neutrophil extracellular trap formation pathway” (Fig 3B). Disease associated pathways like “influenza A”, “epstein barr virus infection”, “pertussis” “salmonella infection” and “kaposi sarcoma associated herpes infection” were also enriched. Enriched terms like “lipid and atherosclerosis”, “pertussis”, “toll-like receptor signalling”, “salmonella infection”, NOD-like receptor signalling pathway”, “Legionellosis” and “neutrophil extracellular trap formation” clustered under legionellosis atherosclerosis extracellular formation (Fig 3D). Clustering of lipid and atherosclerosis pathways with infection associated pathways may indicate association of lipid metabolism with TB outcome. Another cluster was seen for “Kaposi sarcoma associated herpesvirus infection”, “Epstein-Barr virus infection”, “osteoclast differentiation, “C-type lectin receptor signalling pathways, “measles”, “influenza A” and “nectroptosis” under C-type Epstein-Barr differentiation herpesvirus (Fig 3D). The Enrichment map of KEGG pathways shows a number of mutually interconnected pathways with the exception of “antigen processing and presentation” pathway which overlapped with “Epstein-Barr virus infection” pathway (Fig 3F).

The gene concept networks show gene level overlaps between many pathways. For enriched GO Biological Processes top DEG *GBP5,* and other up-regulated genes like *SOCS1* and *PARP14* represent overlap between “response to interferon gamma” and “regulation of response to biotic stimulus” pathways (Supplementary Fig 1A). Top DEGs *AIM2, RSAD2* and *IFIT3* represent overlap between “response to virus”, “defense response to virus” and “defense response to symbiont” pathways. Other top DEGs *CEACAM1* and *SERPING1* are integral to the “regulation of response to biotic stimulus” pathway. For the enriched KEGG pathways, TFs like STAT1, STAT2 and IRF7 overlap between the central pathways indicating commonalities in transcriptional control. Other significant overlaps were present between antigen processing and presentation genes like *TAP1, TAP2, TABP* and *HLA-F* with the Epstein-Barr virus pathway (Supplementary Fig 1B).

### 3. Pathway analysis of down-regulated genes

Enrichment analysis of the 69 down-regulated genes was carried out. Top enriched GO Biological Processes were T cell associated pathways involved with activation and differentiation (Fig 4A). Clustering of enriched terms by tree plots showed clusters such as; *thymic cell selection, apoptotic pathway processes, αβ T cell and mononuclear lymphocyte differentiation, lymphocyte IL4 costimulation production and positive regulation of cell-cell adhesion* (Fig 4C). Enrichment map of the top GO Biological Processes pathways show a high degree of overlap between the enriched pathways (Fig 4D). The top enriched KEGG pathways include the “T cell receptor signalling pathway” overlapping with the enriched GO biological process (Fig 4B). As fewer number of KEGG pathways were enriched for the down-regulated genes, treeplot could not be visualized. The enrichment map for KEGG pathways show overlaps between the “primary immunodeficiency” and both “cell receptor signalling pathway” and “haematopoietic cell lineage” pathways (Figure 4E).

**Figure 4:**
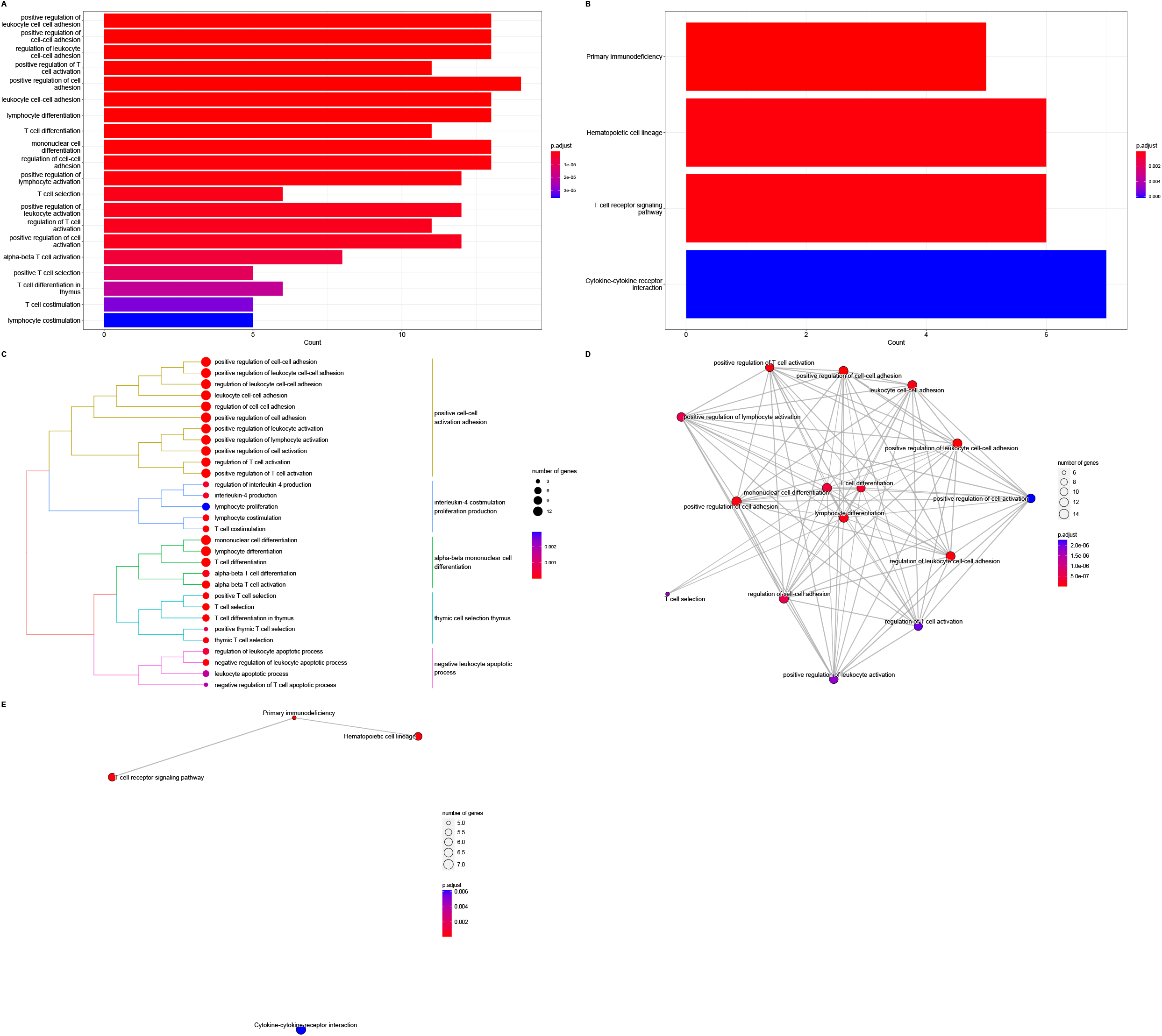
Pathway Analysis for Down-regulated Genes using GO and KEGG pathways: Pathway analysis of GO Biological Processes and KEGG pathways for down-regulated genes show an enrichment of adaptive immune pathways, particularly those associated with T cell activation and differentiation (5A and 5B respectively). Further clustering of these GO Biological Processes pathways show five relevant biological themes associated with adaptive immunity (5C). Enrichment map of GO Biological Processes for down-regulated genes show interconnected pathways (5D) while for KEGG pathways, 3 connected pathways have emerged (5E).

At the gene level, enriched GO Biological Processes have overlap in cell surface CD receptors: CD27, CD6 and *CD3E* (Supplementary Fig 2A). Top DEG, *CCR7* overlaps between the cell adhesion and T cell activation pathways. These pathways also show overlaps in TFs such as *LEF1* and *ETS.* Phorbol ester receptor *PRKCQ,* responsible for activation of AP-1 and NFκB, overlaps between leukocyte cell adhesion pathways. *SKAP1*, a node protein which with its adaptor protein connects cell adhesion with T cell activation (Supplementary Fig 2A). For the enriched KEGG pathways “primary immunodeficiency”, “T cell receptor signalling:” and “hematopoietic cell lineage”, genes *CD3D* and *CD3E* overlap between these pathways. *ICOS* is an overlap between the “T cell receptor signalling” and “primary immunodeficiency” pathway. Other overlaps include *CD40L, IL7R* and *IL11RA* between “primary immunodeficiency”-“cytokine-cytokine receptor interaction”, “primary immunodeficiency”- “hematopoietic cell lineage”-“cytokine-cytokine receptor interaction”, and “hematopoietic cell lineage”-“cytokine-cytokine receptor interaction” respectively (Supplementary Fig 2B).

### 4. Enrichment analysis of DEGs for miRNA targets

miRNAs are important for gene expression regulation. To explore miRNA mediated regulation of these DEGs, ORA was carried out for miRNA targets within the DEG lists. hsa-miR-146a-5p was found to have targets within the up-regulated genes; *PLAUR, NMI, IFITM3, IFITM1, RSAD2, IRF7, FAS, SAMD9L, IFIT3, STAT1, S100A12, TRIM22, IFI44, IFI44L* and *EPSTI1* (Fig 5A). *NMI, PLAUR, IFITM1, IRF7, IFITM3,* and *RSAD2* interacted with hsa-miR-146a-5p and *STAT1* while the rest interacted with hsa-miR-146a-5p alone in the network. Significantly enriched miRNAs targetting the down-regulated genes could not be recovered.

**Figure 5:**
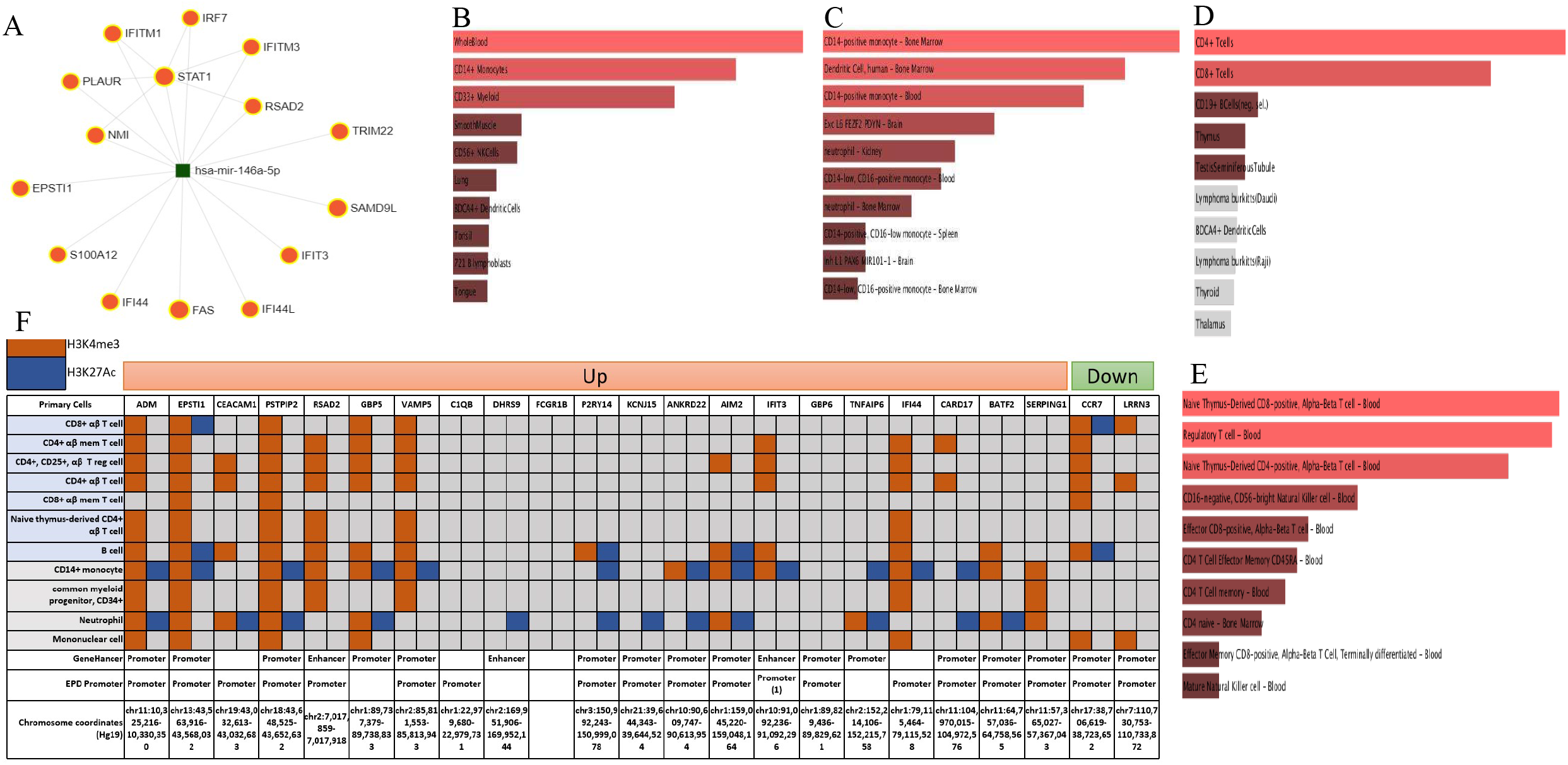
Enrichment of miRNA targets and cell types: The up-regulated genes *NMI, PLAUR, IFITM1, IRF7, IFITM3,* and *RSAD2* interact with hsa-miR-146a-5p and *STAT1* while the rest interacted with hsa-miR-146a-5p alone in the network (5A). Using the Human Gene Atlas and HuBMAP repositories, enrichment analysis was done to identify cell types associated with the up (5B, 5C) and down-regulated genes (5D, 5F) respectively. (5F) Identification of Cell Subtypes: For the top DEGs, presence or absence of histone activation marks (H3K4me3 and H3K27Ac) around the promoters recognized by GeneHancer and EPD promoter database. Presence of histone activatory mark around recognized promoter of a gene indicates the ability of transcriptional activation of the gene in a specific cell type. Here, haematopoietic cells of both lymphoid and myeloid lineage were considered. For each of the top DEGs, the genomic coordinates corresponding to the regulatory region notifications from Genehancer and experimentally validated promoters from EPD were identified. For these regions, the presence or absence of either or both activatory marks were recorded.

### 5. Identification of active cellular subtypes

Considering the 236 up-regulated and 69 down-regulated genes, from Human Gene Atlas and HuBMAP libraries, an enrichment of CD14^+^, CD16^+^, CD33^+^ myeloid cells and dendritic cells was observed for the up-regulated genes (Fig 5B, 5C). This indicates that myeloid cells drive the up-regulated gene signature in response to TB. On the other hand for the down-regulated genes, CD4^+^ and CD8^+^T cell subsets were enriched, concordant with the pathway analysis results (Fig 5D, 5E).

For the top DEGs, ENCODE ChIP-seq experiment data was visualized using UCSC genome browser (hg19) to identify the presence of epigenetic activating marks H3K4me3 and H3K27Ac around regulatory elements characterized by GeneHancer and EPD promoter database in haematopoietic cells (Fig 5F). Up-regulated genes *DHRS*9, *ANKRD22, TNFAIP*6 and *KCNJ*15 have either methylation or acetylation marks around their promoters exclusively in the myeloid lineage cells indicating the capacity of these promoters to initiate the expression of these genes in the myeloid cells. Promoters of genes *ADM, CEACAM*1, *PSTPIP*2, *GBP*5, *VAMP*5, *ANKRD*22, *IFIT*3, *TNFAIP*6 and *IFI*44 contain both tri-methylation and acetlylation peaks in myeloid cells. Genes *ADM, PSTPIP*2, *GBP*5 contained both trimethylation and acetylation peaks in CD14+ monocytes and neutrophils. Genes *CEACAM*1 and *TNFAIP*6 contain both mark in neutrophils alone, while *VAMP*5, *ANKRD*22, *IFIT*3, and *IFI*44 contain both marks in CD14+ monocytes.

*AIM*2 promoter contains both trimethylation and acetylation marks in CD14+ monocytes, neutrophils and B cell. *EPSTI*1 contains both marks in CD14+ monocytes and B cells. While, *PYYR*14 promoter contains both peaks in B cells alone (Fig 5F). This represents an enrichment of both trimethylated and acetylated peaks in myeloid cells indicating formation of active promoters near their TSS in myeloid cells.

Examining the promoters of the two top down-regulated genes, it was observed that *CCR*7 contains both tri-methylated and acetylated promoters in CD8+ αβ T cells and B cells indicating active promoters in these cell types. From the results of enrichment analysis and tragetted visualization it is clear that bulk of the gene expression dysregulation is driven by the myeloid cells including the leading edge of the dysregulation represented by the top up-regulated genes.

### 6. Genetic factors involved in driving the expression of DEGs

GWAS catalog (https://www.ebi.ac.uk/gwas/home) was queried to find GWAS significant SNPs mapped to the top DEGs to identify transcriptomic and genomic overlaps which may help understand disease susceptibility or progression as a function of gene expression. SNPs rs12042284 mapped to *AIM*2 and rs2161037, rs12328794, rs2161037 mapped to *DHRS9* have been reported to be associated with C-reactive protein level. T allele of rs18778419 which is mapped to *C1QB* gene has been observed to be associated with the levels of IFNγ and MCSF. This indicates potential role of IFNγ driving the expression of *C1QB* in tuberculosis.

SNP mapped to *SERPING1*, rs76525968 is reported to be associated with lung function which indicates a role of *SERPING1* in TB disease progression in the lungs in addition to its role in disease diagnosis. SNPs mapped to *AIM2:* rs2518564 and rs2276405 are associated with white blood cell count and psoriasis respectively. Colorectal mucinous adenocarcinoma associated SNP rs1661281 was mapped to *ANKRD22.* Currently, there are no direct overlaps between the genomic loci associated with tuberculosis and these transcriptomic findings. However, few SNPs associated with traits like CRP, IFNγ, MCSF levels map to some of the top DEGs. This may indicate an unexplored role in TB susceptibility.

### 7. Effect of *M.tb* antigens on gene expression in THP1 monocytes and PMA derived macrophages

Among the top 23 DEGs, *AIM*2, *ANKRD*22, *BATF*2, *C*1*QB*, *CARD*17, *CCR*7, *CEACAM*1, *DHRS*9, *EPSTI*1, *FCGR*1*B*, *GBP*5, *LRRN*3 and *SERPING*1 were selected at random and their expression was evaluated in THP1 monocytes and PMA derived THP1 macrophages in the presence or absence of stimulation with *M.tb* antigens Ag85 complex, LAM, CFP10 and ESAT6.

In monocytes, Ag85 complex up-regulated the expression of *AIM2, SERPING1* and *EPSTI1* while significantly down-regulating the expression of *ANKRD22* (Fig 6A, Supplementary Table 1). *EPSTI1* expression was up-regulated by the Ag85 complex in both monocytes and macrophages, however in macrophages a higher fold of up-regulation was observed. Additionally in macrophages, Ag85 complex treatment also significantly up-regulated the expression of *LRRN3, DHRS9* and *FCGR1B* in a descending order of magnitude of expression fold change (Fig 6E Supplementary Table 2). In monocytes, LAM up-regulated the expression of *AIM2, EPSTI1, ANKRD22, SERPING1, GBP5, FCGR1B* and *LRRN3* (Fig 6B, Supplementary Table 1). LAM significantly up-regulated the expression of *EPSTI1, LRRN3* and *GBP5* in macrophages (Fig 6F, Supplementary Table 2).

**Figure 6:**
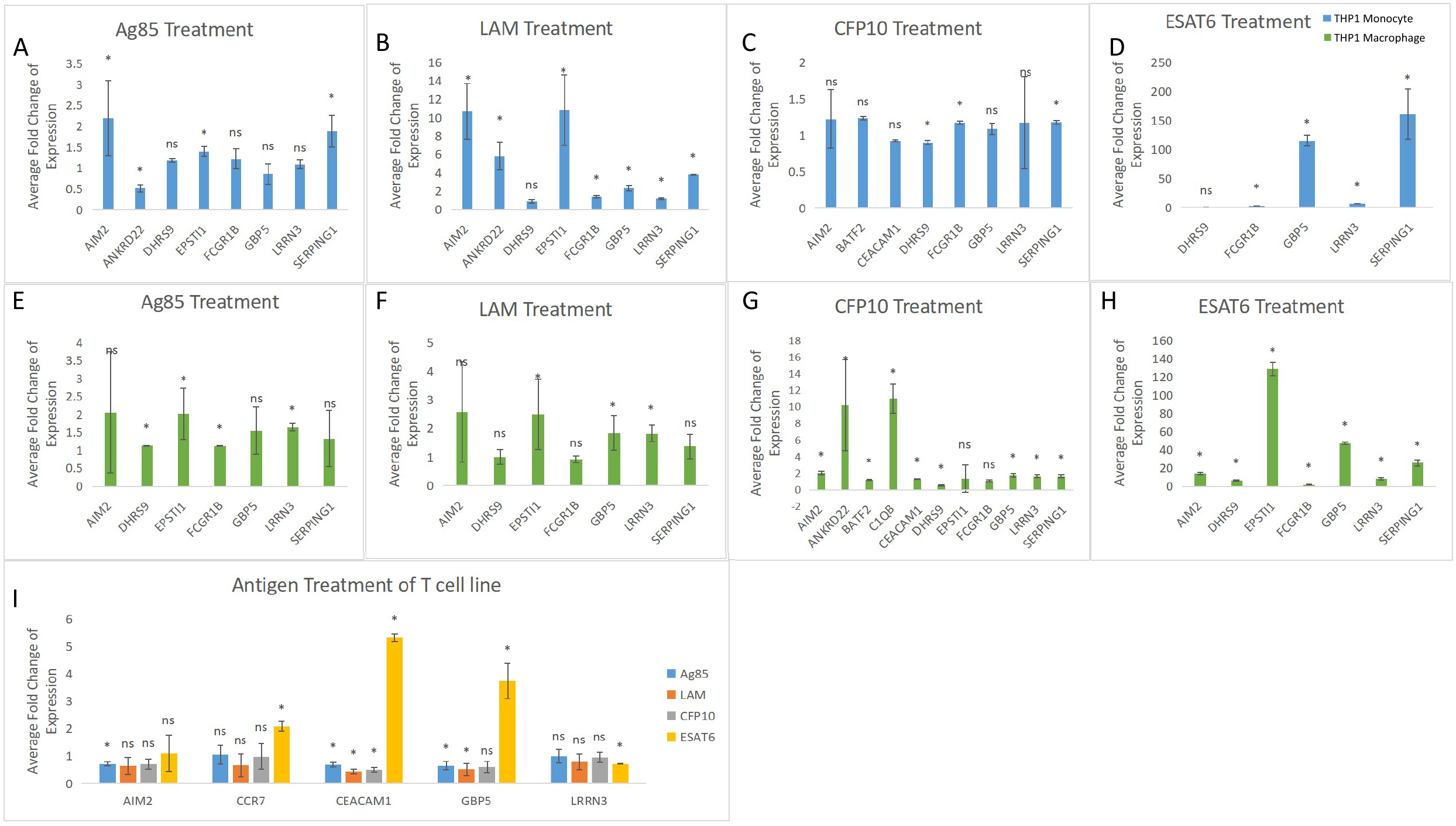
Stimulation of Monocytes and Macrophages with *M.tb* Antigens: Fold change of expression in monocytes after treatment with Ag85 complex, LAM, CFP10 and ESAT6 (6A, 6B, 6C, 6D). Fold change of expression in macrophages after treatment with Ag85 complex, LAM, CFP10 and ESAT6 (6E, 6F, 6G, 6H). Average fold change of expression in T cell line following treatment with Ag85 complex, LAM, CFP10 and ESAT6 (6I); *=p<0.05, ns=not significant

In monocytes, CFP10 induced expression of relatively fewer genes with comparatively lower fold changes of expression than macrophages. It up-regulated the expression of *SERPING1* and *FCGR1B* and mildly down-regulated the expression of *DHRS9* (Fig 6C, Supplementary Table 1). CFP10 up-regulated the expression of *C1QB, ANKRD22, AIM2, GBP5, LRRN3, SERPING1, CEACAM1* and *BATF2* in macrophages (Fig 6G, Supplementary Table 2). Similar to its effect on monocytes, CFP10 also down-regulated the expression of *DHRS9* in macrophages (Fig 6G, Supplementary Table 2). ESAT6 treatment corresponded to the largest magnitudes of expression fold change out of all antigens. In monocytes, ESAT6 showed a comparatively mild up-regulation of *FCGR1B* and *LRRN3* and a much greater fold up-regulation of *SERPING1* and *GBP5* expression (Fig 6D, Supplementary Table 1). In macrophages, ESAT6 up-regulated the expression of *EPSTI1, GBP5, SERPING1, AIM2, LRRN3* and *DHRS9* (Fig 6H, Supplementary Table 2). Similar to monocytes, a lower fold up-regulation of expression was observed for *FCGR1B* in macrophages.

Comparing the performance of *M.tb* antigens in recapitulating the gene signature, in monocytes, fold of up-regulation offered by LAM was distinctly higher compared to the Ag85 complex and CFP10 particularly for genes like, *EPSTI*1, and *AIM*2. ESAT6 treatment had the greatest increase in the fold of expression particularly for *SERPING1* and *GBP*5 in monocytes. In macrophages, ESAT6 treatment showed most up-regulation for the expression of *GBP*5 and *EPSTI*1.

*EPSTI*1 was up-regulated by Ag85 complex, LAM, and ESAT6 in macrophages with ESAT6 inducing the greatest magnitude of up-regulation of gene expression. In monocytes, *EPSTI1* was induced by the Ag85 complex and LAM while protein antigens CFP10 and ESAT6 did not show induction of *EPSTI1* expression. Overall, *EPSTI1* expression was up-regulated by three out of four antigens tested and the effect was seen in either macrophages (ESAT6) or in both monocytes and macrophages (Ag85 complex and LAM) while *SERPING1* was universally upregulated by all of the tested antigens in the monocyte model.

### 8. Effect of *M.tb* antigens on gene expression in Jurkat E6-1 T cells

Stimulation of T cells with the Ag85 complex, LAM and CFP10 down-regulated the expression of *CEACAM1* and *GBP*5, while ESAT6 up-regulated the expression of these genes. ESAT6 also perturbed the expression of the two top down-regulated genes *CCR7* and *LRRN3*, in which the former was up-regulated and the later was down-regulated. Ag85 complex down-regulated the expression of *AIM*2 in T cells (Fig 6I, Supplementary Table 3).

## Discussion

From the publicly available datasets, we identified 305 DEGs in treatment naive pulmonary TB cases with at least 1.2 fold change of expression and FDR corrected p value <0.05. The top DEGs define the leading edge of the gene signature and include 23 genes with greater than 2 fold change of expression. ORA of the up-regulated gene set of 236 genes using GO Biological Processes showed enrichment of IFNγ response pathways within the up-regulated genes: consistent with previous reports [l,3]□. Enrichment of myeloid leukocyte enrichment pathway and other innate immune process pathways observed indicates the importance of the myeloid compartment early in TB.

Top enriched KEGG pathways are involved in pattern recognition like “nod-like receptor signalling”, “c-type lectin receptor signalling pathway” and “toll-like receptor signalling pathway”. “Necroptosis” and “neutrophil extracellular trap formation” pathways were also enriched. Involvement of the TLR pathway in tuberculosis was reported by Maertzdorf et al., 2012 [22]□. CLR Dectin-2, is a receptor of mannosylated LAM which induces T cell responses [23]□. *M.tb* outer membrane protein CpnT induces necroptosis in THP1 derived macrophages through depletion of NAD^+^ [24]□. Formation of NETs has been observed during H37Rv and *M. canetti* infection [25]□. *M.tb* induced NETs stimulate macrophages to produce IL6, TNFα, IL-1β and IL10 [26]□. For the up-regulated genes, most of the enriched pathways has gene-level overlaps in TFs such as *STAT1*, *STAT2* and *IFR7.* Enrichment analysis of the 69 down-regulated genes revealed T cell activation and differentiation pathways for GO biological processes and KEGG pathways. Similarly, for the down-regulated genes TFs engaged in regulation between pathways were *LEF1* and *ETS*.

Exploring their regulation, we performed ORA to enrich miRNAs targetting the DEGs. mir-hsa-146a-5p was observed to target some of the top DEGs like *RSAD2, IFIT3, IFI44,* and *EPSTI1* (Fig 7A). Previously known as miR-146a, this miRNA is transcribed from an independent gene *MIR*146*A.* mir-146a expression is induced by H37Rv and BCG infection of MDMs [27] □. In PBMC, from pulmonary and extrapulmonary TB cases, mir-146a is down-regulated [28,29]□. miR-146a expression inhibits the migration of neutrophils into bronchial epethelium in ashtma [30] □.

GWAS significant SNPs associated with pulmonary TB were not mapped to the top DEGs. SNPs associated with levels of CRP mapped to top DEGs *AIM*2 and *DHRS9* while SNPs associated with IFNγ and MCSF levels mapped to *C*1*QB* indicating a prospective role in driving TB susceptibility. CRP level is elevated in TB independent of HIV status [31]□ and is a biomarker for TB infection. IFNγ activates macrophages in tuberculosis and also helps control replication of *M.tb* bacilli within the macrophages through the production of reactive nitrogen intermediates [32]□. Expression from the *C1QB* gene is generally limited to the monocytes and dendritic cells and its promoter is stimulated by IFNγ [33]□.

Most of the top up-regulated genes were transcriptionally active in the myeloid cells like CD14^+^ monocytes and neutrophils as seen from the accumulation of histone activating marks around their promoters. The bulk of the up-regulated genes were also associated with CD14 myeloid cells. While the down-regulated genes were associated with T cells. This may have resulted from a migration of T cells away from peripheral circulation and into lymph nodes to control the infection.

Immunodominant *M.tb* antigens:Ag85 complex, LAM, CFP10 and ESAT6 were selected to stimulate THP1 monocytes, PMA derived THP1 macrophages and naive T cell line Jurkat E6-1 and study their effect on expression of a subset of the top DEGs. Ag85 complex comprises of fibronectin binding proteins: Ag85A, Ag85B and Ag85C consiting mycolyl-transferase activity which binds mycolipids to arabinogalactan of the mycobacterial cell wall forming the of cord factor. The complex elicits production of Th1 type cytokines. LAM is a mycobacterial cell wall glycolipid. Its mannoside caps are recognized by CLRs on the surface of macrophages, dendritic cells and neutrophils. CFP10 and ESAT6 are coded by genes Rv3874 and Rv3875 on the RD1 region which is associated with virulence. CFP10 is a 10kDa peptide secreted early in mycobacterial lifecycle and has been shown to modulate NO and TNFα secretion from macrophages [34]□. ESAT6 is a mycobaterial virulence factor capable of sequestering β2 microglobulin in the ER with our without its chaperone CFP10, inhibiting antigen presentation. ESAT6 has been shown to stilumate the TLR2/MyD88 pathway evoking a Th17 response [35]□. CFP10, another *M.tb* protein mediates neutrophil activation and migration by regulating intracellular Ca^2+^ levels [36]□.

Compared to the other antigens, ESAT6 induced a greater magnitude of up-regulation of *SERPING1* and *GBP*5 in monocytes and macrophages alike. *SERPING1* was also universally up-regulated by all the tested antigens in the monocyte model, indicating that *SERPING1* up-regulation could be a characteristic of monocytes in TB. GBPs induce expression of *AIM*2 towards activation of AIM2 inflammasome. In our re-analysis, *GBP*5, *GBP*6 and *AIM*2 were identified as top DEGs. Through our stimulation studies, CFP10 and ESAT6 were shown to up-regulate the expression of both *AIM2* and *GBP5* in THP1 macrophages while LAM up-regulated their expression in monocytes. These findings point towards the activation of AIM2 inflammasome early in TB while *M.tb* antigens CFP10, ESAT6 and LAM may induce *AIM2* inflammasome pathway in macrophages and monocytes. In the T cell model, *CEACAM1* was down-regulated significantly by all of the tested antigens while *LRRN3* a top down-regulated gene by ESAT6 treatment.

We also show that *EPSTI1* expression was up-regulated by three of the four antigens, however, most instances of up-regulation was limited to macrophages with the exception of Ag85 complex and LAM treatment where *EPSTI1* expression was up-regulated in both monocytes and macrophages. *EPSTI1* is inducible by *IFNγ* and drives M1 polarization of macrophages through STAT1 and p65 phosphorylation [37]□. STAT1 is another TF which was observed to be up-regulated in whole blood of TB cases. In tuberculosis expression of *EPSTI1* is regulated by hsa-miR-146a-5p which is itself induced by *M.tb* infection.

This study shows that TB specific whole blood transcriptomic signature is defined by up-regulation of innate immune pathways and down-regulation of T cell associated pathways. The bulk and leading edge of the up-regulated gene signature is driven from the myeloid cells (CD14+ monocytes and neutrophils) while the bulk of the down-regulated signature is driven CD4+ and CD8+ T cells as observed from both ORA and targetted visualization. The induction of the leading edge genes or the top DEGs was observed in a myeloid cell model of THP1 monocytes and macrophages. Overall, *SERPING1* overexpression was induced by all antigens in the monocytes while *EPSTI1* expression was induced primarily in macrophages. In the T cell model, genes *CEACAM1* was down-regulated by all antigens expect ESAT6. However, ESAT6 significantly down-regulated the expression of top down-regulated gene *LRRN3* thereby replicating the down-regulated gene signature.

## 4. Acknowledgements

The authors would like to thank the National Institute of Biomedical Genomics (NIBMG, Kalyani) for carrying out this work. Anuradha Gautam would like to thank UGC, GOI for fellowship support

## 6. Authorship Contributions

**Conceptualization:**BP

**Data curation:** AG, SKM, BP

**Formal analysis:** AG SKM BP

**Investigation:** BP, AG, SKM

**Methodology:** AG, SKM, BP

**Resources:** BP

**Supervision:** BP, SKM

**Validation:** BP, AG

**Visualization:** AG, BP

**Writing – original draft:** AG, BP

**Writing – review & editing:** AG, BP, SKM

## 9. Abbreviations

DEGs: Differentially Expressed Genes
CFP10: Culture Filtrate Protein 10
ESAT6: Early Secreted Antigenic Target 6kDa
LAM: Lipoarabinomannan
Ag85: Antigen 85 Complex
LTBI: Latent TB Infection
PLWH: People Living With HIV
Xpert: MTB/RIF
TST: Tuberculin Skin Test
ATT: anti-tuberculosis therapy
GWAS: Genome-wide Association Study
CRP: C-reactive Protein
GEO: Gene Expression Omnibus
FDR: False Discovery Rate
ORA: Over Representation Analysis

---

Supplementary Figure 1: **Gene Level View of Overlaps Between Pathways Enriched from Up-Regulated Genes:** The gene concept network shows the overlaps between genes among the GO biological processes enriched pathways (1A) and KEGG pathways (1B).

Supplementary Figure 2: **Gene Level View of Overlaps Between Pathways Enriched from Down-Regulated Genes:** The gene concept network shows the overlaps between genes among the GO biological processes enriched pathways (2A) and KEGG pathways (2B).

