## Supplementary material for "Major Contribution of Myeloid Cells In TB specific Host Gene Signature: Revelations from Re-Analysis of Publicly Available Datasets": Supple Fig1

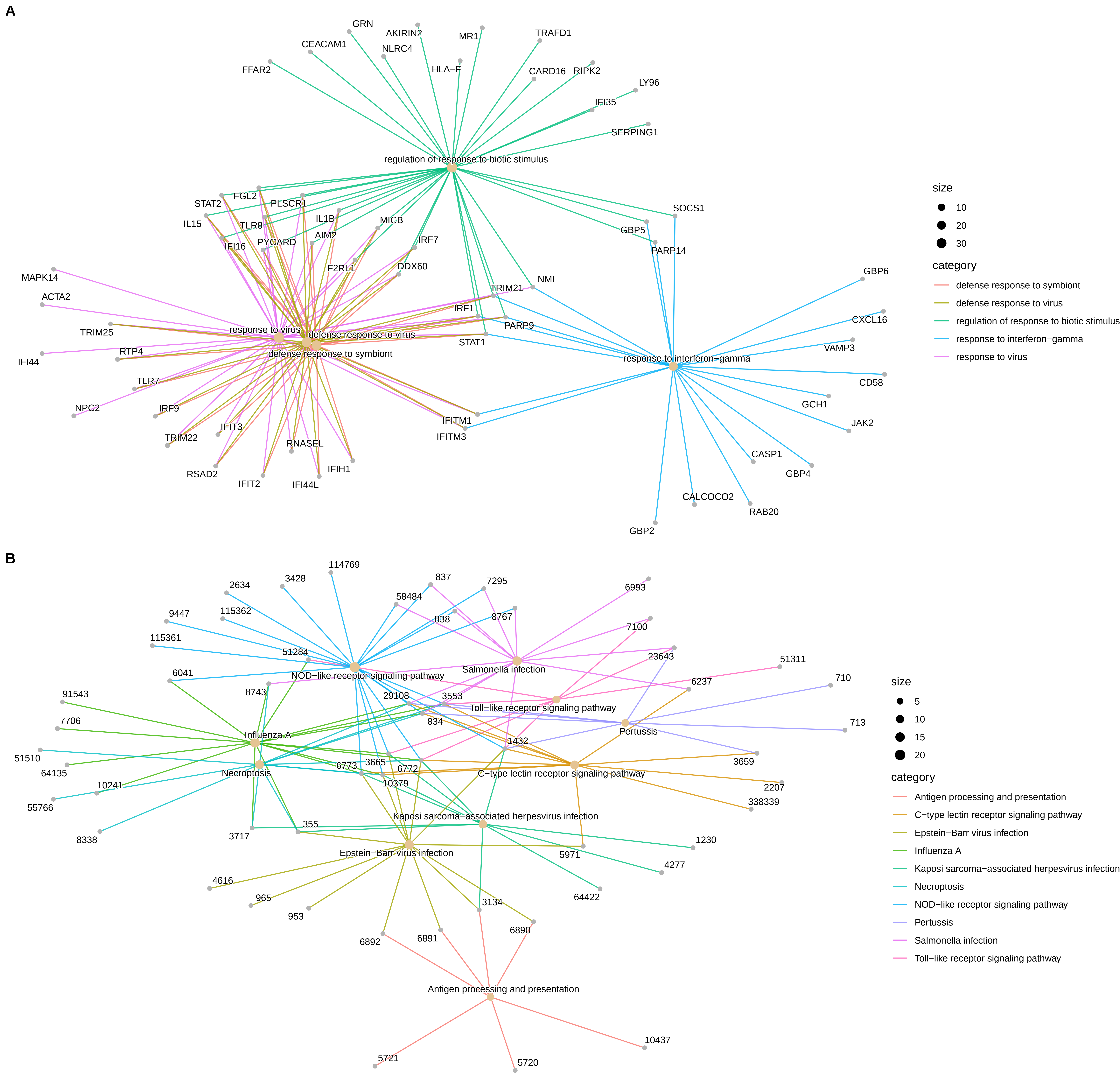


Supplementary Figure 1: Gene Level View of Overlaps Between Pathways Enriched from Up−Regulated Genes: The gene concept network shows the overlaps between genes among the GO Biological Processes enriched pathways (1A) and KEGG pathways (1B).
