## Supplementary material for "Major Contribution of Myeloid Cells In TB specific Host Gene Signature: Revelations from Re-Analysis of Publicly Available Datasets": Supple Fig2

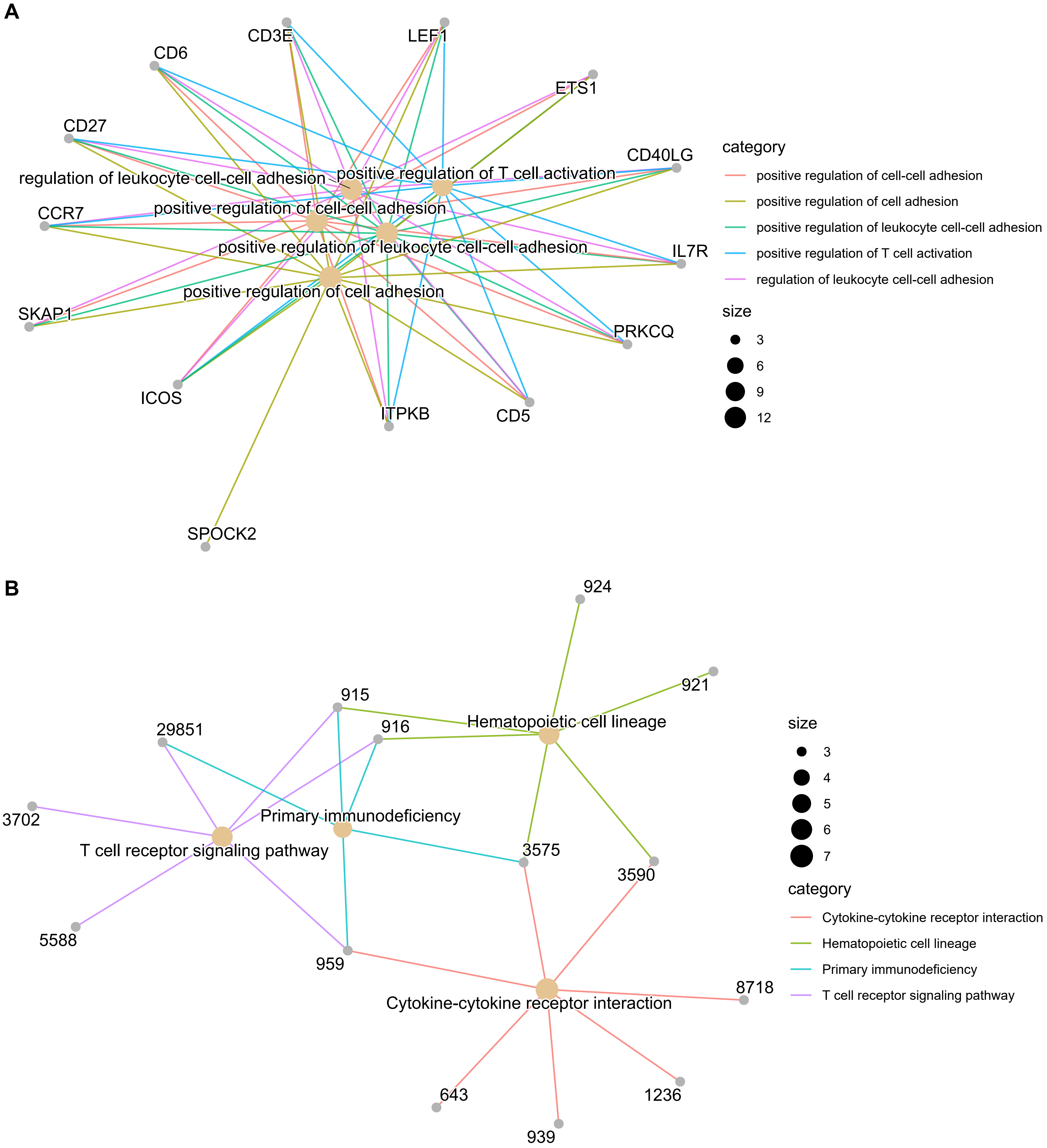


Supplementary Figure 2: Gene Level View of Overlaps Between Pathways Enriched from Down-Regulated Genes: The gene concept network shows the overlaps between genes among the GO Biological Processes enriched pathways (2A) and KEGG pathways (2B).
