## Supplementary material for "Major Contribution of Myeloid Cells In TB specific Host Gene Signature: Revelations from Re-Analysis of Publicly Available Datasets": Supple Table 1,2,3

Supplementary Table 1: Expression of Top DEGs following antigen stimulation in THP1 monocytes

| Gene | THP1 MONOCYTE | | | | | | | |
|---|---|---|---|---|---|---|---|---|
| | Ag85 Complex | | LAM | | CFP10 | | ESAT6 | |
| | Average Fold Change | p value | Average Fold Change | p value | Average Fold Change | p value | Average Fold Change | p value |
| **AIM2** | 2.193 | 0.025 | 10.718 | 0.001 | 1.222 | 0.597 | NA | NA |
| **ANKRD22** | 0.515 | 0.006 | 5.816 | 0.005 | NA | NA | NA | NA |
| **BATF2** | NA | NA | NA | NA | 1.236 | 0.078 | NA | NA |
| **C1QB** | NA | NA | NA | NA | NA | NA | NA | NA |
| **CARD17** | NA | NA | NA | NA | NA | NA | NA | NA |
| **CCR7** | NA | NA | NA | NA | NA | NA | NA | NA |
| **CEACAM1** | NA | NA | NA | NA | 0.927 | 0.644 | NA | NA |
| **DHRS9** | 1.188 | 0.073 | 0.908 | 0.368 | 0.901 | 0.027 | 0.990 | 0.94 |
| **EPSTI1** | 1.397 | 0.009 | 10.830 | 0.002 | NA | NA | NA | NA |
| **FCGR1B** | 1.217 | 0.111 | 1.397 | 0.048 | 1.173 | 0.010 | 3.155 | 0.006481136 |
| **GBP5** | 0.860 | 0.250 | 2.326 | 0.025 | 1.088 | 0.284 | 115.161 | 7.25524E-05 |
| **LRRN3** | 1.091 | 0.224 | 1.201 | 0.024 | 1.173 | 0.758 | 7.123 | 0.001342007 |
| **SERPING1** | 1.882 | 0.006 | 3.791 | 0.013 | 1.179 | 0.007 | 160.789 | 0.000133507 |

Supplementary Table 2: Expression of Top DEGs following antigen stimulation in PMA derived THP1 Macrophages

| Gene | PMA Derived THP1 Macrophage | | | | | | | |
|---|---|---|---|---|---|---|---|---|
| | Ag85 Complex | | LAM | | CFP10 | | ESAT6 | |
| | Average Fold Change | p value | Average Fold Change | p value | Average Fold Change | p value | Average Fold Change | p value |
| **AIM2** | 2.060 | 0.332 | 2.579 | 0.120 | 2.005 | 0.001 | 14.115 | 6.33011E-05 |
| **ANKRD22** | NA | NA | NA | NA | 10.186 | 0.009 | NA | NA |
| **BATF2** | NA | NA | NA | NA | 1.203 | 0.020 | NA | NA |
| **C1QB** | NA | NA | NA | NA | 10.979 | 0.00005 | NA | NA |
| **CARD17** | NA | NA | NA | NA | NA | NA | NA | NA |
| **CCR7** | NA | NA | NA | NA | NA | NA | NA | NA |
| **CEACAM1** | NA | NA | NA | NA | 1.290 | 0.031 | NA | NA |
| **DHRS9** | 1.129 | 0.022 | 0.999 | 0.902 | 0.498 | 0.002 | 6.627 | 0.000101112 |
| **EPSTI1** | 2.017 | 0.022 | 2.503 | 0.031 | 1.339 | 0.636 | 128.769 | 1.821E-06 |
| **FCGR1B** | 1.121 | 0.003 | 0.929 | 0.319 | 1.094 | 0.075 | 1.868 | 0.00769399 |
| **GBP5** | 1.548 | 0.119 | 1.846 | 0.026 | 1.765 | 0.003 | 47.103 | 3.46163E-06 |
| **LRRN3** | 1.643 | 0.003 | 1.830 | 0.004 | 1.635 | 0.004 | 8.369 | 4.82141E-05 |
| **SERPING1** | 1.326 | 0.520 | 1.376 | 0.109 | 1.632 | 0.002 | 25.738 | 1.11762E-05 |

Supplementary Table 3: Expression of Top DEGs following Antigen treatments in Jurkat E6-1 T cell line

| Gene | JURKAT E6-1 T CELL | | | | | | | |
|---|---|---|---|---|---|---|---|---|
| | Ag85 Complex | | LAM | | CFP10 | | ESAT6 | |
| | Average Fold Change | p value | Average Fold Change | p value | Average Fold Change | p value | Average Fold Change | p value |
| **AIM2** | 0.7192 | 0.0454 | 0.6454 | 0.1272 | 0.7130 | 0.1607 | 1.1109 | 0.9823 |
| **ANKRD22** | NA | NA | NA | NA | NA | NA | NA | NA |
| **BATF2** | NA | NA | NA | NA | NA | NA | NA | NA |
| **C1QB** | NA | NA | NA | NA | NA | NA | NA | NA |
| **CARD17** | NA | NA | NA | NA | NA | NA | NA | NA |
| **CCR7** | 1.0523 | 0.8680 | 0.6684 | 0.1564 | 0.9885 | 0.7554 | 2.0890 | 0.00028 |
| **CEACAM1** | 0.6874 | 0.0030 | 0.4382 | 0.0001 | 0.4986 | 0.0342 | 5.3249 | 0.00004 |
| **DHRS9** | NA | NA | NA | NA | NA | NA | NA | NA |
| **EPSTI1** | NA | NA | NA | NA | NA | NA | NA | NA |
| **FCGR1B** | NA | NA | NA | NA | NA | NA | NA | NA |
| **GBP5** | 0.6551 | 0.0359 | 0.5179 | 0.0417 | 0.6077 | 0.0552 | 3.7409 | 0.00098 |
| **LRRN3** | 0.9962 | 0.8680 | 0.7874 | 0.1793 | 0.9632 | 0.6561 | 0.7183 | 0.00052 |
| **SERPING1** | NA | NA | NA | NA | NA | NA | NA | NA |

Table 4: Primer sequences used for the RTPCR experiments

| S.No | Gene | Sense | Primer Sequence | No. of Bases | UCSC *In silico* PCR | Tm (℃) | Size of Amplicon | PMID |
|---|---|---|---|---|---|---|---|---|
| 1 | AIM2 | F | TCGGCACAGTGGTTTCTTAG | 20 | >uc001ftj.1__AIM2:884+1020 | 58.9 | 137bp | 31836304 |
| | AIM2 | R | GCTGAGTTTGAAGCGTGTTG | 20 | | 59.6 | | |
| 2 | ANKRD22 | F | AAAGCACTGACAGGTGTTTGCTA | 23 | >uc001kfj.4__ANKRD22:205+401 | 61.2 | 197bp | |
| | ANKRD22 | R | TTGGCAGATGGGCTCAGAGT | 20 | | 63.3 | | |
| 3 | BATF2 | F | AGACCCCAAGGAGCAACA | 18 | >uc001ocf.1__BATF2:170+308 | 59.2 | 139bp | 25762344 |
| | BATF2 | R | CAGGGCGAGGTTGTCTTT | 18 | | 58.8 | | |
| 4 | C1QB | F | AAGGTGCCCGGTCTCTACTA | 20 | >uc001bgd.3__C1QB:619+752 | 58.8 | 134bp | 21698244 |
| | C1QB | R | ACCTGGAAGGTGTTGTAGGC | 20 | | 59.1 | | |
| 5 | CARD17 | F | GGACTCTCAGCAGGTCCAAC | 20 | >uc001pir.1__CARD17:279+410 | 59.8 | 132bp | |
| | CARD17 | R | AACTCTTTCAGTGCTGGGCAT | 21 | | 61.2 | | |
| 6 | CCR7 | F | GTCTTCGGTGTCCACTTTTGC | 21 | >uc002huw.3__CCR7:442+541 | 62 | 100bp | 25359998 |
| | CCR7 | R | CGTAGCGGTCAATGCTGATG | 20 | | 62.7 | | |
| 7 | CEACAM1 | F | CAGTCACCTTGAATGTCACCTATG | 24 | >uc002oub.3__CEACAM1:818+962 | 61.01 | 145bp | |
| | CEACAM1 | R | GTTCCATTGATAAGCCAGGAGTAC | 24 | | 61.01 | | |
| 8 | DHRS9 | F | TGTGCTTCTGGATGTGACCG | 20 | uc010zdc.2__DHRS9:529+618 | 63.3 | 90bp | |
| | DHRS9 | R | AGACCCCAGAGACCTTTCTCC | 21 | | 61 | | |
| 9 | EPSTI1 | F | CCCGCAATAGAGTGGTGAACT | 21 | >uc001uyx.2__EPSTI1:114+306 | 61.4 | 193bp | |
| | EPSTI1 | R | TATGCACTTGTGCGCCTCTG | 20 | | 63.5 | | |
| 10 | FCGR1B | F | GGAAGGGGTGCACCGGAAGG | 20 | >uc001eip.3__FCGR1B:854+951 | 70.3 | 98bp | 20861863 |
| | FCGR1B | R | CACGGGGAGCAAGTGGGCAG | 20 | | 70.6 | | |
| 11 | GBP5 | F | GAGAATTCCCTAAGGCCAAAGC | 22 | >uc001dnc.3__GBP5:1138+1312 | 62.5 | 175bp | |
| | GBP5 | R | CAGGCTCTAGCTCATCATCAGGC | 24 | | 64.8 | | |

| S.No | Gene | Sense | Primer Sequence | No. of Bases | UCSC *In silico* PCR | Tm (℃) | Size of Amplicon | PMID |
|---|---|---|---|---|---|---|---|---|
| 12 | LRRN3 | F | TGGTACCATTGAGTCTCTGCCA | 22 | uc022akc.1__LRRN3:1053+1226 | 62.9 | 174bp | 31199560 |
| | LRRN3 | R | TGCCGAACATTCTGACCTTGG | 21 | | 65.6 | | |
| 13 | SERPING1 | F | CCAAGATGCTATTCGTTGAACCC | 23 | >uc010rjv.1__SERPING1:358+435 | 63.9 | 78bp | 26511510 |
| | SERPING1 | R | TGGTGGCTGAATTGGTTGTTG | 21 | | 64.1 | | |
| 14 | B-ACTIN | F | CATGGATGATGATATCGCCGC | 21 | >uc003sos.2__ACTB:36+141 | 63.7 | 106bp | |
| | B-ACTIN | R | CACGATGGAGGGGAAGACG | 19 | | 64 | | |
| 15 | GAPDH | F | ACAACTTTGGTATCGTGGAAGG | 22 | >uc001qop.2__GAPDH:671+771_ | 63.8 | 101bp | |
| | GAPDH | R | GCCATCACGCCACAGTTTC | 19 | | 66.8 | | |

F= Forward
R=Reverse
